# Assessment of local potato cultivars found in *Cis-Himalayan* region of West Bengal through morphology and biochemical profiling

**DOI:** 10.1101/2022.04.27.489635

**Authors:** Subir Dutta, Subhra Chakraborty, Bodeddula Jayasankar Reddy, Sumita Nag, Sahanob Nath, Sanghamitra Routh, Vivekananda Behera, Gnanasing Jesumaharaja Lazar, Birudukota Monika, Lakshmi Hijam, Moumita Chakraborty, Suvendu Kumar Roy, Ashok Choudhury, Satyajit Hembram, Manoj Kanti Debnath, Rupsanatan Mandal

**Affiliations:** Department of Genetics and Plant Breeding, Uttar Banga Krishi Vishwavidyalaya, Pundibari, Coochbehar, West Bengal, India; Department of Plant Pathology, Uttar Banga Krishi Vishwavidyalaya, Pundibari, Coochbehar, West Bengal, India; Department of Biotechnology, Ramakrishna Mission Vivekananda Education and Research Institute, Howrah, West Bengal, India; Department of Genetics and Plant Breeding, Centurion University of Technology, Paralakhemundi, Gajapati, Odisha, India; Department of Soil Science, Uttar Banga Krishi Vishwavidyalaya, Pundibari, Coochbehar, West Bengal, India; Department of Agriculture Statistics, Uttar Banga Krishi Vishwavidyalaya, Pundibari, Coochbehar, West Bengal, India; Regional Research Station, Terai Zone, Uttar Banga Krishi Vishwavidyalaya, Pundibari, Coochbehar, West Bengal, India

**Keywords:** Potato, North Bengal, Diabetes, DUS characterization, Anthocyanin, Glycemic index

## Abstract

Potato is a major global food crop grown for tubers (underground storage stems) that are high in carbohydrates, minerals, and vitamins. The presence of anthocyanins in tuber skin and flesh can have a significant impact on customer preferences. Potatoes are also high in resistant starches, which have a variety of health benefits, including enhanced fullness, cholesterol reduction, and a reduced risk of colon cancer etc. In West Bengal, diabetes is prevalent. Similarly, according to ICMR studies, colon cancer affects 8.9% of all cancer patients in West Bengal cancer which is caused by ill eating habits or the consumption of foods that are low in antioxidants. 5-10% of colon cancers are genetically caused, while the remainder are caused by poor eating habits or the consumption of foods that are low in antioxidants. To address these issues, one strategy is to eat foods with a low glycemic index and high antioxidant content as a staple food. The tuber tissues of the potato (*S. tuberosum* L.) accumulate various quantities of anthocyanins, which are commonly consumed around the world. Anthocyanins are pigments that range in colour from red to purple and are found throughout the plant kingdom. Anthocyanins are powerful antioxidants that are water soluble. The red skin potato is well-known among the general public. The epidermal layer contains a high quantity of red anthocyanins, which causes the skin to be red. With this background the present study has been undertaken to address the following objectives morphological (quantitative and qualitative traits) and biochemical characterization of local potato cultivars and identification of most stable genotypes based on the anthocyanin content and tuber yield of local potato cultivars. For our recent research 14 local potato cultivar from North Bengal were collected and evaluated for DUS characterization showed wide range of variability with respect to different phenotypic variants of ten characters. Overall predominant DUS characters of local potato cultivars found in North Bengal are medium sized apical length of sprout, short stem height, compact foliage structure, open leaf structure, ovate lanceolate type leaflet shape, purple leaf sprout predominant colour, spherical leaf sprout shape, medium intensity of anthicyanin coloraton at the base, light intensity of anthicyanin coloraton at the tip and weak nature of light sprout pubescence base. The results of the present investigation suggest that local potato cultivars collected from the northern part of West Bengal showed a high level of genetic variation. The differences between genotypes were highly significant at the 1% and 5% alpha level for all characters like tuber yield, length and weight of tuber, length of sprout, chlorophyll content, canopy temperature, and number of tuber per plant, according to the analysis of variance. Three quantitative traits namely tuber yield, tuber breadth, tuber length played major role in the genetic variance. Anthocyanin content had more contribution to diversify the local potato cultures according to biochemical characterization AMMI analysis suggested that Jalpai from CoochBehar is the most stable potato cultivars in respect to anthocyanin content and tuber yield per plant.

## Introduction

Potato (*Solanum tuberosum* L.) from family solanaceae is the main non-cereal food consumed worldwide (1) and the vegetable with the highest antioxidant contribution to human diet (2). As of nutritive value, raw potato contains 26% carbohydrate, 2% dietary fiber and 4% protein (USDA). Along with all these potatoes also have high antioxidant and are a good source of several vitamins like vitamin B complex and minerals like K, Mg and Fe. Various potatoes with pigmented skin contain carotenoids like anthocyanin and phenolic acids (3, 4, 5, 6). Anthocyanins are health-promoting compounds found in purple and red tuber skin/flesh potato cultivars. They are high in antioxidants, support gut bacteria, and lower the glycemic index of starch (7,8). Methylation, acylation, and polyglycosilation are examples of post-biosynthetic changes that extend the types of anthocyanins generated and may affect their bioactive qualities (9). The potato produces the most anthocyanins with the highest acylation levels among the Solanaceae plants. During food technological processes, acylation with organic acids (typically caffeic, p-coumaric, and ferulic acids) improves the physical chemical stability of anthocyanins (10, 11). Beside this anthocyanins are flavonoids that can be found in a variety of foods, primarily fruits and vegetables. Because of the conjugated bond in their structures, they absorb light in the 500 nm spectrum. Fruits and vegetables have vibrant red, blue, and purple colours (12, 13,14). Because of their antioxidant and anti-inflammatory qualities, anthocyanins have been shown in epidemiologic studies to reduce the incidence of heart and circulatory illness, diabetes, arthritis, and even cancer (15). These incidences can be reduced by eating foods high in anthocyanins. However, the vast majority of Indians, who are primarily poor, are unable to obtain high-priced anthocyanin-rich crop items or fruits.

Several factors determine the phytochemical content of potato tubers. Each potato variety has its own nutritional qualities, which are determined by its genetic history. Environmental–agronomic factors, as well as their interplay with genotype, influence the amount and kind of health-promoting chemicals naturally accumulating in tubers (16, 5).

Local potatoes are currently threatened by genetic erosion as a result of the introduction of high-yielding variants. Characterizing the local genetic resources for crop development in emerging nations like India is a vital and important task (17,18). Local potato cultivars are useful breeding materials for successful hybridization to increase sustainable potato production (19,20). The genetic makeup of such native potatoes located in West Bengal’s northern region has yet to be investigated. The basic stage in increasing production is determining the genetic relatedness of local potatoes (21). According to the recommendations issued (22), indigenous potato cultivars must always meet the criteria of distinctness, homogeneity, and stability. DUS testing is one of the most essential criteria for determining the distinctness, uniformity, and stability of local cultivars (23). Morhological and phenome research, which are based on the measurement of a plant’s vegetative and reproductive structure, are crucial for the identification and use of local cultivars (24,25).

## Material & methods

### Plant material

Fourteen local potato genotypes were collected and maintained by the Central Germplasm Conservation Unit (CGCU), Directorate of Research, Uttar Banga Krishi Viswavidyalaya, Cooch Behar, and West Bengal, India. Before being used as plant materials in the current studies, selected potato genotypes were evaluated for five years by CGCU, UBKV (**Table 1)** by Reddy et al. 2021. The whole tuber of the fourteen potato cultivars were collected from the field after harvesting for the biochemical characterization. Potatoes were collected when they were fully mature and delivered to the lab right away. But, anthocyanin content was estimated only for whole tuber for the samples bring from the Kanksha, West Burdwan and Narendrapur, Kolkata.

**Table 1.** Details of the plant materials using the present study.

| Sl. No. | Cultivars | CGCU Entry No | Population type |
| --- | --- | --- | --- |
| 1 | Cooch Behar Deshi 01 | CGCU/2015/Potato/16 | RSWF |
| 2 | Cooch Behar Deshi 02 | CGCU/2015/Potato/17 | RSWF |
| 3 | Jalpaiguri Local 01 | CGCU/2015/Potato/18 | RSWF |
| 4 | Cooch Behar Local 01 | CGCU/2015/Potato/19 | WSWF |
| 5 | Cooch Behar Jalpai 01 | CGCU/2015/Potato/20 | RSRF |
| 6 | Cooch Behar Local 02 | CGCU/2015/Potato/21 | WSWF |
| 7 | K. Sundari | CGCU/2015/Potato/22 | RSWF |
| 8 | Jalpaiguri Local 02 | CGCU/2015/Potato/23 | WSWF |
| 9 | Cooch Behar Badami 01 | CGCU/2015/Potato/24 | RSRF |
| 10 | Cooch Behar Badami 02 | CGCU/2015/Potato/25 | RSWF |
| 11 | K. Jyoti | CGCU/2015/Potato/26 | WSWF |
| 12 | Cooch Behar Local 03 | CGCU/2015/Potato/27 | RSWF |
| 13 | Jalpaiguri Local 03 | CGCU/2015/Potato/28 | WSWF |
| 14 | Cooch Behar Local 04 | CGCU/2015/Potato/29 | RSWF |

### Data collection and statistical analysis

Most of the qualitative and quantitative traits were observed on individual plant basis. Morphological data were recorded on five randomly selected plants by excluding the plants located at borders to avoid border effect so that highest precision could be achieved. For qualitative features, only the first replication of the experiment was evaluated, but all replications were included for quantitative aspects. The descriptions and instructions provided by the PPV & FR Authority were used to capture qualitative characteristics. The qualitative characteristics were analysed using Microsoft Office Excel 2007 and NTSYSpc v. 2.2 (26) software. GENRES v. 3.11 (27) and SPSS v. 16.0 software were used to analyse quantitative data (28). **(Table 2)**.

**Table 2.** Qualitative and quantitative trait for characterization of potato genotypes.

| *Quantitative traits* |  | *Qualitative traits* |  |
| --- | --- | --- | --- |
| Traits | Instructions | Traits | Plant parts |
| Length of apical sprout | On the 30th day after release from cold storage, the length of the apical sprout (cm) was measured from the base to the tip with a Vernier Caliper. | Predominant colour; Shape; Intensity of Anthocyanin colouration at base of sprout; Intensity of Anthocyanin colouration at sprout tip; Pubescence base; Length of apical sprout | Light sprout |
| Germination % | Germination % = (Number of plantlets emerged per plot )/(Total number of tuber planted per plot ) × 100 | Foliage structure; Height of main stem (cm) | Whole plant |
| Plant height (PH) | Plant height was measured using a measuring scale from the ground level to the tip of the plant 50 days after planting, during the period of maximum leaf development. | Solidity; Cross section | Stem |
| Leaf length and width | Leaf length and width was observed on 4th fully developed leaf from the top of the plant by measuring scale. | Structure | Leaf |
| Number of tubers per plant and weight per plant | Total number of tubers from randomly selected five plants were counted and weight is determined | Lateral leaflet shape | Leaflet |
| Tuber length and width | With a Vernier Caliper, the length and width of 10 marketable tubers from each plot was measured. | Predominant skin Colour; Secondary skin colour; Predominant colour of flesh<br>Secondary colour of flesh | Tuber |

### Stability Analysis

Potato tuber yields and anthocyanin content were measured in three different Indian locales. The genotype-environment interaction (GEI) reduces the phenotypic and genotypic values’ correlation. The genotype adaptation and stability index (GEI) can be investigated to understand more about genotype adaptation and stability. The genotype-environment interaction (GEI) reduces the connection between phenotypic and genotypic values, which causes gene effect estimates to be skewed. Combining abilities for particular attributes that are subject to environmental changes is a common theme in GEI research. A typical multivariate approach is additive main effects and multiplicative interaction (AMMI) analysis.

The AMMI stability value (ASV) described by Purchase et al. (2000) was calculated as follows:

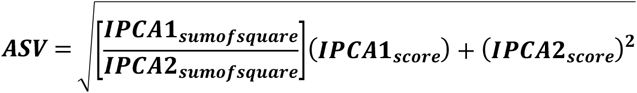

The weight assigned to the IPCA1 value by dividing the IPCA1 sum of squares by the IPCA2 sum of squares is SSIPCA1/SSIPCA2. The greater the IPCA score, whether negative or positive, the better a genotype is suited to certain settings. Lower ASV scores suggest a genotype that is more stable in different contexts.

Total anthocyanin content estimation

Total anthocyanin extraction was carried out using the technique described by Leite-Legatti et al., 2012 (30), with minor modifications. Each cultivar’s whole tuber peel, ski, and flesh (0.5 g of each type of sample) were placed in a test tube with 15 mL of extract solution (95 percent ethanol: 1.5 mol/L hydrochloric acid = 85:15, v/v), ultrasonically mashed and crushed for 30 minutes (45°C, 100 W), and centrifuged at 5000g for 5 minutes. After collecting the supernatant, the precipitate was removed once again. Two supernatants were mixed and preserved in the refrigerator at -20°C for analysis.

The anthocyanin concentration was calculated using the following formula (31, 32, 30), which states that anthocyanin has a maximum absorption peak at pH 1 and no absorption peak at pH 4.5 under 525 nm. For 15 minutes in the dark, the anthocyanin extract was combined with 25 mM hydrochloric acid-potassium chloride buffer (pH 1) or 0.4 M sodium acetate buffer (pH 4.5). After that, the mixes’ optical density was measured at 700 nm and 525 nm, and the anthocyanin content was determined using the formula (Equations (1) and (2)). The absorbance ratio A420/A525 of anthocyanin was also measured to be utilised as a deterioration index (DI) since anthocyanin would change to chalcone in an unstable environment.

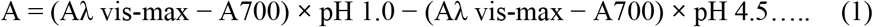

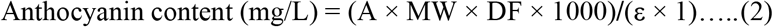

(The molecular weight of anthocyanin was calculated as cyanidin-3-glucoside (449.2), DF: dilution factor, E: molar absorption coefficient (ε) was calculated as cyanidin-3-glucoside (26900), and 1: quartz cuvette diameter (cm)).

### Resistant starch estimation and glycaemic index estimation

The RS and GI kit (Megazyme International Ireland Ltd.) was used to estimate the RS concentration and GI in potatoes. The RS content and GI was determined using a 510 nm laser and expressed as a percentage of dry matter (33).

## Result and Discussion

### DUS characterization-

#### Frequency distribution of phenotypic variants

The potato cultivars evaluated for DUS characterization showed wide range of variability with respect to different phenotypic variants of ten characters.

Overall predominant DUS characters of local potato cultivars found in North Bengal are medium sized apical length of sprout, short stem height, compact foliage structure, open leaf structure, ovate lanceolate type leaflet shape, purple leaf sprout predominant colour, spherical leaf sprout shape, medium intensity of anthicyanin coloraton at the base, light intensity of anthicyanin coloraton at the tip and weak nature of light sprout pubescence base. **Rest characters are showed in Table 3**.

**Table 3.** DUS characterization basis on quality.

| Sr. No. | Genotype % | Character | Quality |
| --- | --- | --- | --- |
| 1. | 85.14 | Stem height | Short |
| 2. | 14.29 |  | Medium |
| 3. | 50 | Foliage structure | Compact |
| 4. | 28.57 |  | Semi Compact |
| 5. | 21.43 |  | Open |
| 6. | 28.57 | Leaf sprout predominant colour | White |
| 7. | 7.14 |  | Pink |
| 8. | 14.29 |  | Red purple |
| 9. | 42.86 |  | Purple |

**(Detailed DUS characterization are described in our previous investigation by Reddy et al. 2021.)**

#### Phenotypic allelic diversity and ANOVA

The polymorphism percentage, average number of distinct allele (Na), average number of effective allele (Ne), Nei’s gene diversity (h), and Shannon’s information index were used to determine phenotypic allelic diversity from the three groups (RSRF, RSWF, and WSWF) of potato genotypes (I). The RSRF group had 0% polymorphism, the RSWF group had 79.31 percent polymorphism, and the WSWF group had 89.66 percent polymorphism, for a total of 56.32 percent polymorphism. The RSRF group had the lowest Na, Ne, and lowest value for I, h, and uh, whereas the WSWF group had the highest Na, Ne, and highest value for I,h, and uh, but the RSWF group had a medium value for all parameters. The aggregate total potato genotypes in this study had values of 1.264, 1.352, 0.311, 0.208, and 0.252 for the Na, Ne, I, h, and uh parameters, respectively **(Estimated different allelic diversity parameters based on 29 phenotypic allele of ten DUS traits given in supplementary Table 1)**. The results of analysis of variance (ANOVA) showed that 98% variation within the population and 2% variation among the population (groups). The differences between genotypes were highly significant at the 1% and 5% alpha level for all characters like tuber yield, length and weight of tuber, length of sprout, chlorophyll content, canopy temperature, and number of tuber per plant, according to the analysis of variance. The tuber length varies from 1.61 to 7.13 c.m. Number of tubers per plant ranges from 2 to 37. Also the tuber yield differs significantly across varieties and ranges from 0.02 to 0.54 kg. **(Table 2 ANOVA and genetic components analysis based on nine quantitative traits of potato given as supplementary file)**. These data substantiate the incidence of variations among the varieties in a significant manner. **(Analysis of variance among the three groups of potato revealed by DUS traits given in supplementary Table 3)**.

#### Cluster analysis based on DUS traits

As a result of dendrogram analysis using the UPGMA method, all fourteen potato cultivars were divided into four clusters with a similarity coefficient of 0.60. **(Detailed cluster analysis given in Figure 1 as supplementary file)**. Cluster I and Cluster IV had four genotypes each, followed by Cluster II and Cluster III, which had three genotypes each. The genetic similarity of the cultivars ranged from 37.93 percent (Cooch Behar Badami 01 and Jalpaiguri Local 03) to 100 percent (Cooch Behar Badami 01 and Jalpaiguri Local 03). (Cooch Behar Deshi 01 and Cooch Behar Deshi 02).

**Figure 1:**
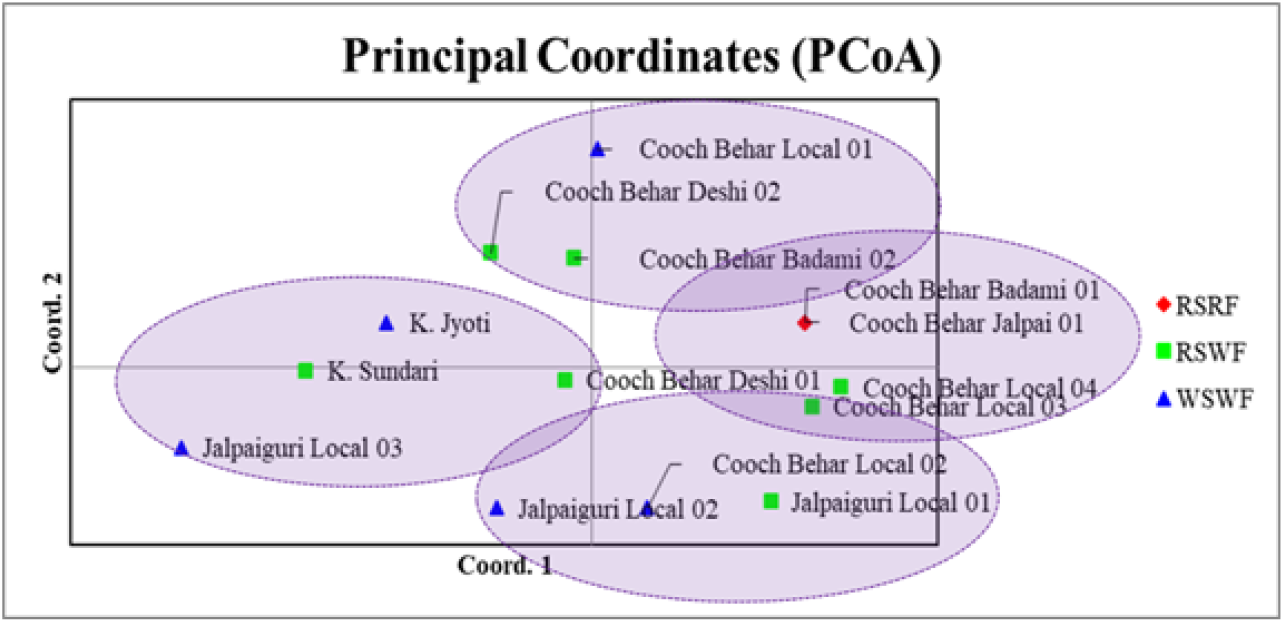
PCoA plot accomplished using 29 phenotypic variants of ten DUS traits

#### Principal coordinates analysis (PCoA)

According to the PCoA data, the first three coordinates account for 55.87 percent of total variation. Overall, the results showed that genotypes from the WSWF group had the most variation within the population. Allelic diversity and ANOVA analysis corroborated these findings. According to PCoA, all of the genotypes clustered into four groups, comparable to dendrogram analysis **(Figure 1)**.

In the present experiment, three groups of potato (RSRF, RSWF and WSWF) were considered based on their skin and flesh colour. Skin colour was determined to be red in maximum potato genotypes, but groups WSWF were white or yellow. As such, the gene pool is considered a valuable initial resource for plant breeding, because it contains co-adapted gene complexes with tolerance or adaptation to diseases and specific ecological conditions, and many plant species (34,35). To be useful for plant breeders, genetic resources must be characterized by morphological and agronomic traits (36,37, 38). For this reason, there is need to collect, characterize and evaluate remnant local genotypes before they disappear (39, 40, 41). The main goal of potato breeding is to develop potential varieties that ensure the highest and stable production in a range of environments (41,42, 43). The cluster analysis has different genotypes on the basis of similarity and thus provides a hierarchical classification (44). The clustering of local potato genotypes, collected from Artvin and Rize provinces in the Eastern Black Sea Region of northern Turkey, on a dendrogram into twenty-seven separate groups resulted from their different morphological and agronomic characteristics. The present research has identified the relationship between local potato genotypes. In general, no association was observed for clusters within the collection zone. This lack of association may be a result of tuber transport from village to village or from province to province by humans. Another possibility, may be that it suited the ecological conditions of the region, and these local genotypes are widely accepted by the farmers for reasons such as cooking quality and taste of tubers (45, 46). Several differences among pinto bean populations (47), sweet potato populations (48,49), potato genotypes (50,51,52,46) and winter squash genotypes (51) were observed for most of the morphological and agronomic characters, as shown in the present study. Racho et al. (2001) (52) stated that 113 wild potato clones were clustered into 51 subgroups in terms of isoenzymatic patterns, whereas Karuri et al. (2010) (53) found that dendrogram was highly variable among the morphological characters of 89 sweet potato genotypes and as such, obtained phenotypic characters that separated the genotypes into two major clusters. It is important to select the lines that are superior in terms of genetic diversity and agronomical properties during the improvement studies (54,55). Karaca (2004) (56) selected 9 genotypes among 63 potato genotypes in terms of maturity time, tuber shape and plant height. However, Escribano et al. (1997) (57) and Galvan et al. (2006) (58) have suggested that the use of morphological traits should be complemented with more accurate techniques to achieve reliable evaluation and characterization of species diversity. Therefore, in recent years, successful results could be obtained using DNA markers and molecular techniques in the determination of genetic traits for variety improvement (47,50). Besides, conservation and maintenance of this valuable genetic material is necessary, because these populations are an important diversity source which could be used in breeding programmes (59). An analysis of morphological and agronomic traits showed that genetic variation was high among the local potato genotypes sampled. Clusters obtained in this study may provide a basis for further study, and it could be selected separately for each in terms of traits investigated further as potato breeding programs. These evaluations could assist breeders to select and identify genotypes with desirable characteristics for inclusions in variety breeding programs.

#### Phenotypic and genotypic coefficient of variation

Higher magnitude of PCV (phenotypic coefficient of variation) and GCV (genotypic coefficient of variation) (> 20%) were observed for sprout length at 30 days from the cold storage (29.74 and 38.79%), chlorophyll content (25.62 and 24 %), tuber length (32.36 and 29.07 %), total tubers per plant (108.39 and 100.43 %), breadth of tuber (38.51 and 28.86 %), tuber yield per plot (74.29 and 86.21%). But low PCV and GCV was observed for canopy temperature and internodal length at 90 DAP (20.80 %).

This reveals that influence of the environmental factors on these characters is negligible for characters like chlorophyll content and canopy temperature, in case of other traits environment plays moderate role in expressing the phenotypes. The complete manifestation of the phenotype is determined by genotypic performance. Sharma (2004) (60), Basavaraj et al., (2005) (61), and Engida et al., (2006) (62) all came to similar conclusions.

#### Heritability

It is difficult to establish the relative quantity of heritable and non-heritable components of variation present in the population using only the genotypic coefficient of variation. Estimates of heritability and genetic advance would supplement this parameter. Estimate heritability was recorded high for the characters viz., chlorophyll content (87%), number of tubers per plant (85%), tuber length (80%), tuber yield (74%) and tuber breadth (68%). The moderate heritability was perceived for sprout length (58%) and canopy temperature (53%).

The high to moderate heritability recorded for the traits indicated that these characters were less influenced by environmental fluctuations and governed by the additive gene effects that are substantially contributing towards the expression of these traits. Hence, selection for these traits will lead to accumulation of more desirable genotypes. The present findings on heritability are in accordance with findings reported by the various workers viz., 63, 64,65.

#### Genotypic and phenotypic correlation coefficients

In the present investigation, genotypic and phenotypic correlations among seven quantitative characters were estimated to study how tuber yield is influenced by its component characters. The approximations of genotypic and phenotypic correlation coefficients have been represented in **(Table 4 Genotypic (G) and Phenotypic (P) correlation for seven quantitative characters in Potato in supplementary file)**.

**Table 4.** ANOVA and genetic components for seven biochemical properties in potato.

| SOV | d.f. | AWT | ASK | AFL | GIF | GIC | RSF | RSC |
| --- | --- | --- | --- | --- | --- | --- | --- | --- |
|  |  | Mean of sum square |  |  |  |  |  |  |
| Replication | 2 | 49.7 | 137 | 0.62 | 5.44 | 97 | 234.178 | 0.08 |
| Genotype | 13 | 9963.1** | 66850** | 146.21** | 9.19** | 61.78** | 18.26** | 2.79** |
| Error | 26 | 2.6 | 1 | 0.304 | 0.67 | 0.23 | 0.38 | 0.01 |
| Max |  | 174.35 | 479.5 | 21.18 | 29.06 | 93.21 | 56.66 | 3.05 |
| Min |  | 5.36 | 53.4 | 0 | 21 | 73.9 | 40 | 0.14 |
| Coefficient variation % | ECV | 2.7 | 0.61 | 20.09 | 3.36 | 0.57 | 1.31 | 9.73 |
|  | GCV | 96.89 | 85.44 | 253.96 | 6.88 | 5.41 | 5.18 | 70.44 |
|  | PCV | 96.93 | 85.45 | 254.75 | 7.66 | 5.44 | 5.34 | 71.11 |
| Heritability |  | 0.99 | 0.99 | 0.99 | 0.8 | 0.98 | 0.93 | 0.98 |

“\*” significance at 5% probability; “\*\*” significance at 1% probability
[AWT: Anthocyanin content in whole tuber; ASK: Anthocyanin content in skin; AFL: Anthocyanin content in flesh; GIF: Glycaemic index of fresh potato; GIC: Glycaemic index of cooked potato; RSF: Resistant starch in fresh potato; RSC: Resistant starch in cooked potato]

In general, the genotypic and phenotypic correlation exhibited similar trend but in most of the cases genotypic correlation were at higher magnitude than phenotypic correlations. Very close values of genotypic and phenotypic correlation were also observed between some character’s combinations, which might be due to reduction in error (environmental) variance to minor proportions as reported by (66). Patel et al., (2018) (67) also conveyed that lower phenotypic correlation than genotypic correlation represents an inherent association amid numerous characters.

Tuber yield was found positively correlated with chlorophyll content, tuber length, tuber breadth, and number of tuber per plant at both genotypic and phenotypic levels indicating the importance of these characters for determining tuber yield.

While choosing characters that have a direct impact on yield, associations with other characters should be taken into account as well, as this will have an indirect impact on yield. Positive and significant correlation at both phenotypic and genotypic levels were observed in case of length of sprout with chlorophyll content and canopy temperature; chlorophyll content with length of sprout, number of tubers per plant and tuber yield; canopy temperature with length of sprout and tuber length; tuber breadth with tuber length; tuber length with canopy temperature, tuber breadth and tuber yield; tuber breadth with canopy temperature, tuber length and tuber yield; number of tubers per plant with chlorophyll content and tuber yield.

Significant negative correlations in the experiment were observed for tuber yield with length of sprout and canopy temperature at genotypic level indicating negative influence of this character on tuber yield. Also canopy temperature showed negative significant association with number of tubers per plant at genotypic level. In general, length of sprout revealed negative correlation with maximum characters. Pleiotropy and/or linkage may also be the genetic reasons for this type of negative association. The pleotropic genes that affected both characters in the desired direction will be strongly acted upon by selecting and rapidly brought towards fixation.

The result of correlation coefficient implied that the number of tubers per plant, tuber yield and chlorophyll content may be considered for selection towards yield improvement from the population of potato in our present study.

#### Cluster analysis

A dendrogram **(Figure 2 dendrogram based on seven quantitative traits given in supplementary file)** constructed using UPGMA method, which was grouped all fourteen potato cultivars into six clusters based on seven morphological quantitative traits. Cluster I consisted with four number of genotypes followed by cluster III and cluster IV which included three number of genotypes respectively. Cluster II and Cluster VI included only one genotypes respectively. Cluster V have the two gnotypes. Interesting fact that, cluster I have the genotypes with higher anthocyanin content. However, the grouping based on morphological quantitative traits does not meet the grouping based on their pigmentation in skin and flesh.

**Figure 2.**
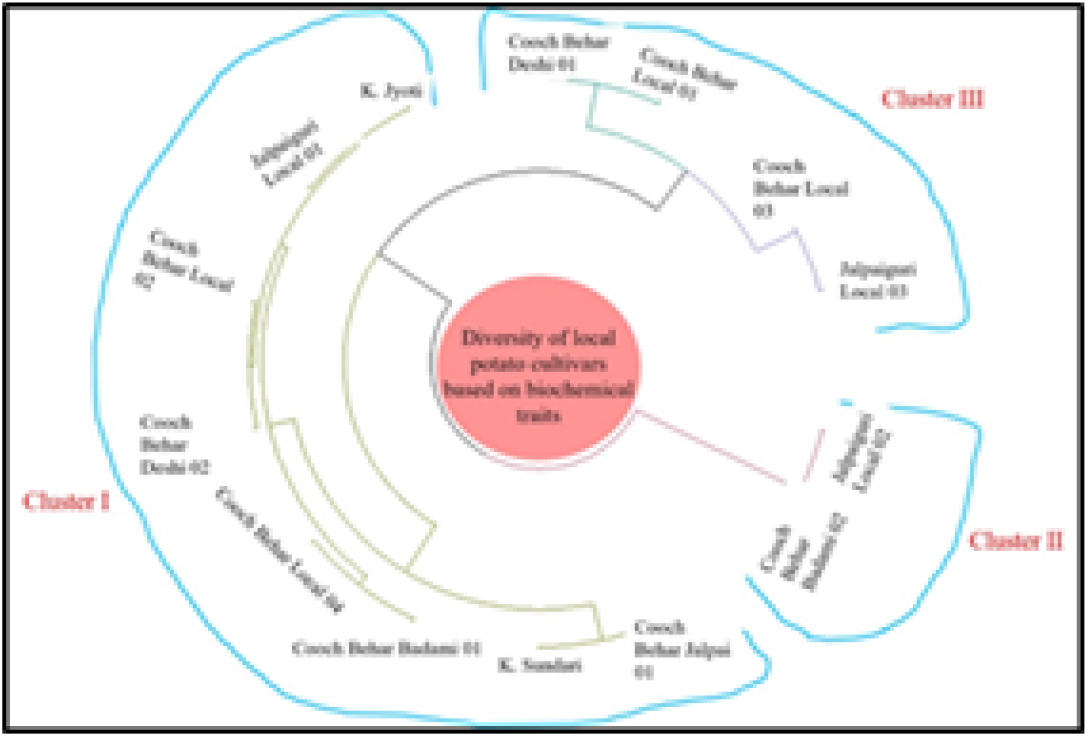
Dendrogram based on seven biochemical properties

### Biochemical characterization

#### Analysis of variance (ANOVA)

All the fourteen genotypes showed significant variation based on their seven biochemical properties. The differences between genotypes were highly significant at the 1% and 5% alpha level for all characters like anthocyanin content (whole tuber, skin and flesh), glycaemic index (fresh potato and cooked potato) and resistant starch (fresh potato and cooked potato) according to the analysis of variance **(Table 4)**.

#### Phenotypic and genotypic coefficient of variation

Higher magnitude of PCV and GCV (> 20%) were experiential for anthocyanin content (whole tuber) (97% and 97%), anthocyanin content (skin) (85%), anthocyanin content (flesh) (254 and 255%) and resistant starch (cooked) (71 and 70%). Lower magnitude of GCV and PCV were observed for glycaemic index (fresh) (7 and 8%), glycaemic index (cooked) (5%), and resistant starch (fresh) (5%).

Therefore, additive component is predominant in case of higher GCV and PCV observations. This reveals that influence of the environment for these characters is negligible in the full expression of the phenotype. Whereas the characters showing lower PCV and GCV are influenced by environmental factors in larger aspect but there is no such traits in this experiment that showed environmental influence.

#### Heritability

It is difficult to establish the relative quantity of heritable and non-heritable components of variation present in the population using only the genotypic coefficient of variation. Estimates of heritability and genetic progress might be used to round out this number. For all of the characteristics under consideration, the heritability estimate was high (>95%).

The high and moderate heritability recorded for the traits indicated that these characters were less influenced by environmental fluctuations and governed by the additive gene effects that are substantially contributing towards the expression of these traits. Hence, selection for these traits will lead to accumulation of more desirable genotypes.

#### Genotypic and phenotypic correlation coefficients

In the present investigation, genotypic and phenotypic correlations among twelve quantitative characters were estimated to study how tuber yield is influenced by its component characters. The estimates of genotype and phenotype correlation coefficients have been presented in **Table 5**.

**Table 5.**
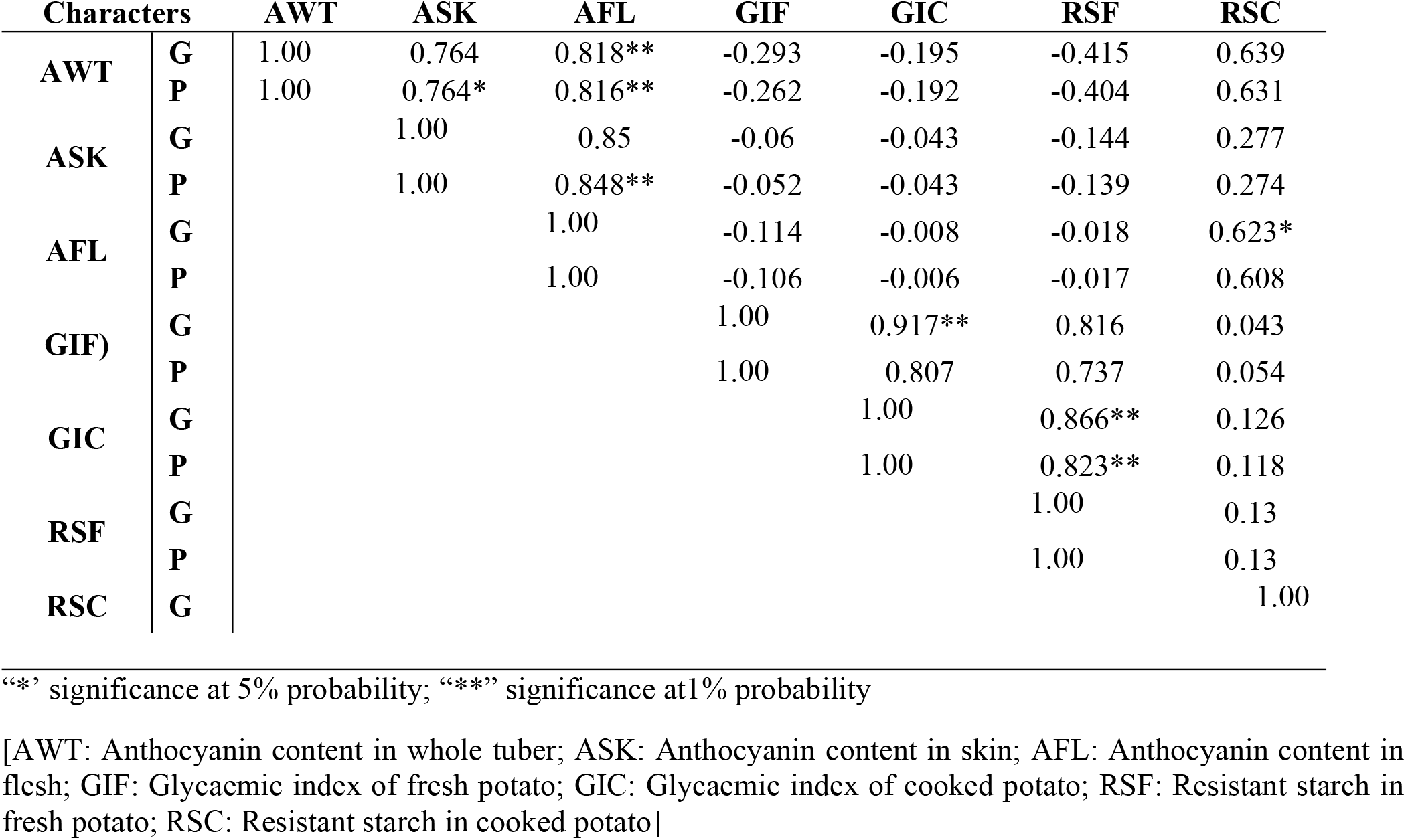
Genotypic (G) and Phenotypic (P) correlation for seven biochemical properties in Potato.

| Characters |  | AWT | ASK | AFL | GIF | GIC | RSF | RSC |
| --- | --- | --- | --- | --- | --- | --- | --- | --- |
| AWT | G | 1.00 | 0.764 | 0.818** | -0.293 | -0.195 | -0.415 | 0.639 |
|  | P | 1.00 | 0.764* | 0.816** | -0.262 | -0.192 | -0.404 | 0.631 |
| ASK | G |  | 1.00 | 0.85 | -0.06 | -0.043 | -0.144 | 0.277 |
|  | P |  | 1.00 | 0.848** | -0.052 | -0.043 | -0.139 | 0.274 |
| AFL | G |  |  | 1.00 | -0.114 | -0.008 | -0.018 | 0.623* |
|  | P |  |  | 1.00 | -0.106 | -0.006 | -0.017 | 0.608 |
| GIF) | G |  |  |  | 1.00 | 0.917** | 0.816 | 0.043 |
|  | P |  |  |  | 1.00 | 0.807 | 0.737 | 0.054 |
| GIC | G |  |  |  |  | 1.00 | 0.866** | 0.126 |
|  | P |  |  |  |  | 1.00 | 0.823** | 0.118 |
| RSF | G |  |  |  |  |  | 1.00 | 0.13 |
|  | P |  |  |  |  |  | 1.00 | 0.13 |
| RSC | G |  |  |  |  |  |  | 1.00 |

“\*” significance at 5% probability; “\*\*” significance at 1% probability
[AWT: Anthocyanin content in whole tuber; ASK: Anthocyanin content in skin; AFL: Anthocyanin content in flesh; GIF: Glycaemic index of fresh potato; GIC: Glycaemic index of cooked potato; RSF: Resistant starch in fresh potato; RSC: Resistant starch in cooked potato]

Three quantitative traits namely anthocyanin content (skin), anthocyanin content (flesh), anthocyanin content (whole tuber). Played major role in the genetic variance followed by resistant starch (cooked) and glycaemic index (cooked). Around seven principal components were obtained. Three traits viz., anthocyanin content (skin), anthocyanin content (flesh), anthocyanin content (whole tuber) were contributed more for PC1. Whereas Glycaemic Index (fresh), glycaemic index (cooked), resistant starch (fresh) were contributed much for PC2.

But, glycaemic Index (fresh), glycaemic index (cooked), resistant starch (fresh) had negative association with PC1, while rest of the traits positively associated with PC1. But all these traits of PC1 do not give negative effect in PC2. Detachment of each variable with respects to PCl and PC2 showed the influence of this variable in the discrepancy of germplasm **(Figure 2)**.

The result of correlation coefficient implied that anthocyanin content (skin); anthocyanin content (flesh) and resistant starch (cooked) may be considered for selection for quality improvement in the population of potato under study.

#### Principal component analysis

Principal component analysis was used to determine the role of several biochemical variables in phenotypic diversity. As demonstrated in **Table 6**, it produced Eigen values and percent of variance for seven main component axes across 14 potato cultivars. It was discovered that the first two main components together accounted for 82.65% of the overall variance across all cultivars studied. Combination of the first, second and the third principal components accounted for 93.06% of the aggregate phenotypic variation among the Seven component axes of the all the potato cultivars. Among the seven PCs, Eigen values for PC1, PC2 and PC3 were 3.22, 2.55 and 0.72 respectively. Scree plot clarified the fraction variance concomitant with each principal component obtained by drawing a graph between eigen values and Principal component numbers.

**Table 6.** Summary of the PCA for biochemical properties.

| Principal Component | Eigen value | Variance (%) | Cumulative variation (%) |
| --- | --- | --- | --- |
| PC1 | 3.229 | 46.129 | 46.129 |
| PC 2 | 2.558 | 36.546 | 82.675 |
| PC 3 | 0.727 | 10.391 | 93.066 |
| PC 4 | 0.307 | 4.387 | 97.453 |
| PC 5 | 0.129 | 1.850 | 99.303 |
| PC 6 | 0.041 | 0.591 | 99.893 |

Three quantitative traits namely anthocyanin content (skin), anthocyanin content (flesh), anthocyanin content (whole tuber). Played major role in the genetic variance followed by resistant starch (cooked) and glycaemic index (cooked). Around seven principal components were obtained. Three traits viz., anthocyanin content (skin), anthocyanin content (flesh), anthocyanin content (whole tuber) were contributed more for PC1. Whereas Glycaemic Index (fresh), glycaemic index (cooked), resistant starch (fresh) were contributed much for PC2.

But, glycaemic Index (fresh), glycaemic index (cooked), resistant starch (fresh) had negative association with PC1, while rest of the traits positively associated with PC1. But all these traits of PC1 do not give negative effect in PC2. Detachment of each variable with respects to PCl and PC2 showed the influence of this variable in the discrepancy of germplasm **(Figure 3 contribution of seven biochemical traits towards variability using PCA given as supplementary file)**.

**Figure 3.**
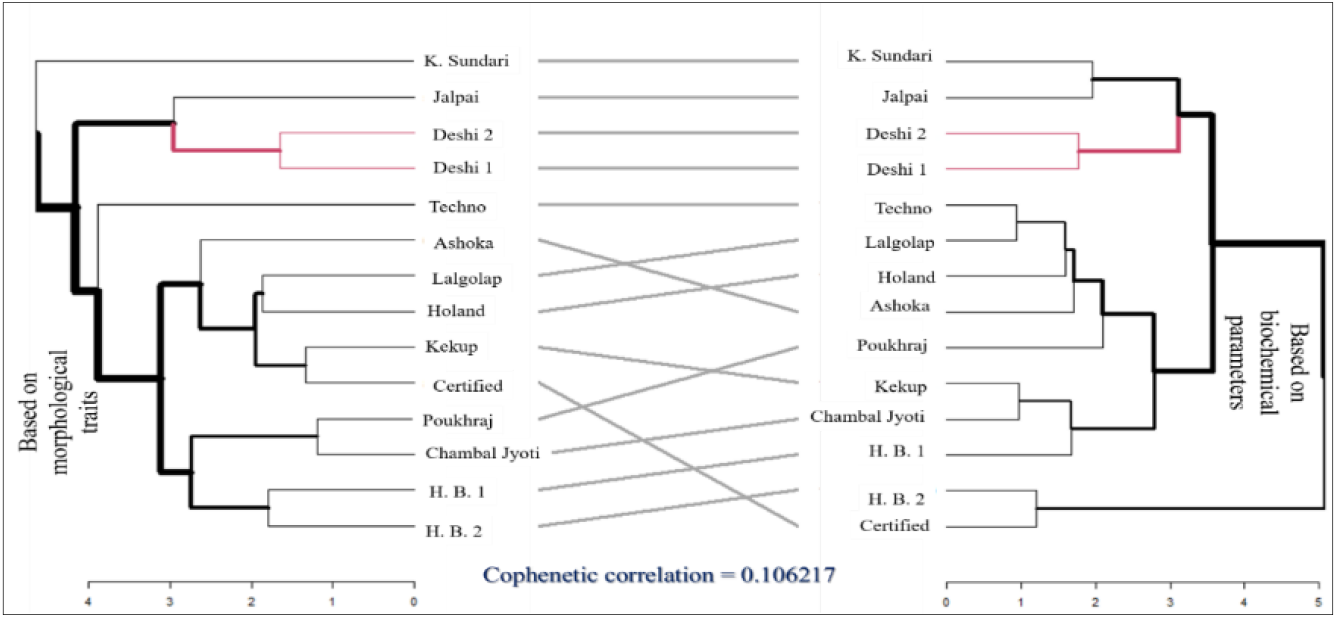
Comparative analysis between dendrogram based on biochemical and dendrogam based on quantitative traits

#### Cluster analysis

A dendrogram **(Figure 2)** constructed using UPGMA method, which was grouped all fourteen potato cultivars into three clusters based on seven biochemical traits. Cluster I had eight genotypes whereas cluster-II had 2 genotypes, cluster III had four genotypes.

#### Comparative analysis among the quantitative, qualitative and biochemical charaterization

In present investigation, dendrogram generated based on qualtative character quantitative character and biochemical character were likened with the help of co-phenetic relations. Details of the co-phenetic relationship value among the dendrogram were represented by various figures.

The results specify that relationship between dendrogram based on biochemical and dendrogram based on quantitative traits were positive and non-significant **(Figure 3)**.

Similarly, relation between dendrogram based on quantitative traits and dendrogram based on qualtative characters were found negative and non-significant **(Figure 4)**.

**Figure 4.**
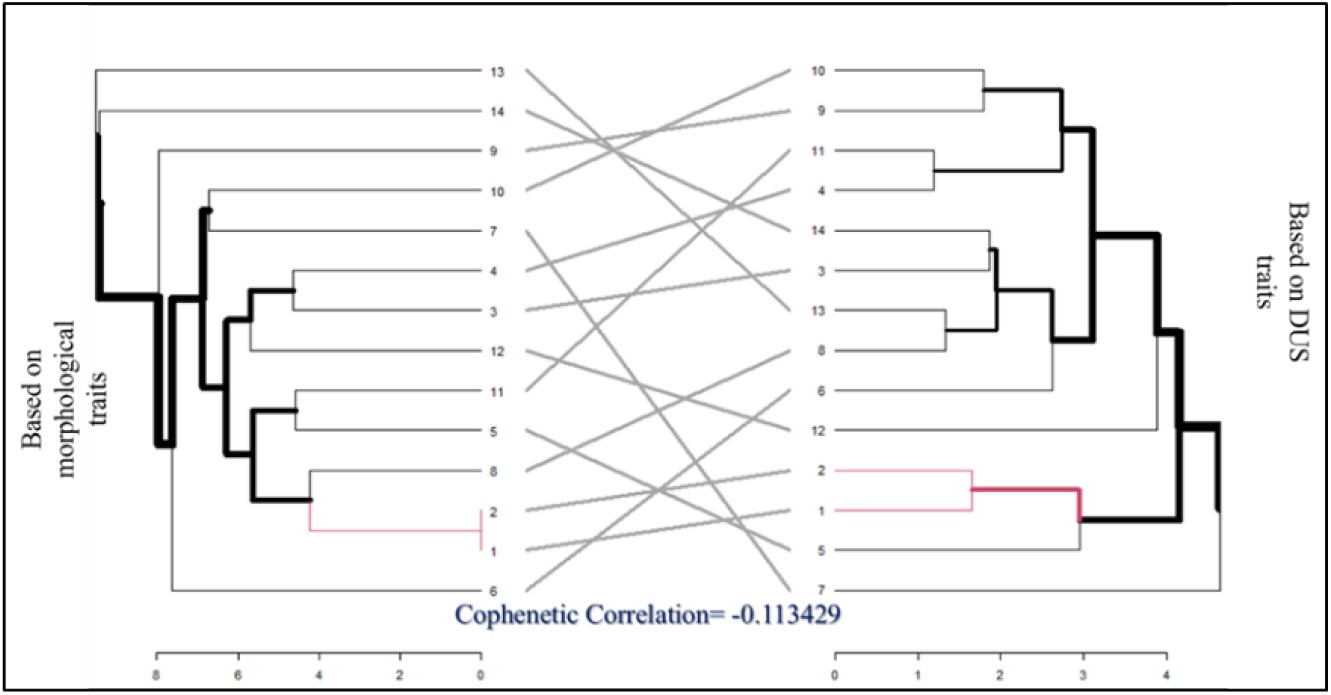
Comparative analysis between dendrogram based on qualitative and dendrogam based on quantitative traits.

Results achieved from comparing dendrogram based on qualtative characters and dendrogram based on biochemical characters showed that there was negative and non-significant relationship between them **(Figure 5)**.

**Figure 5.**
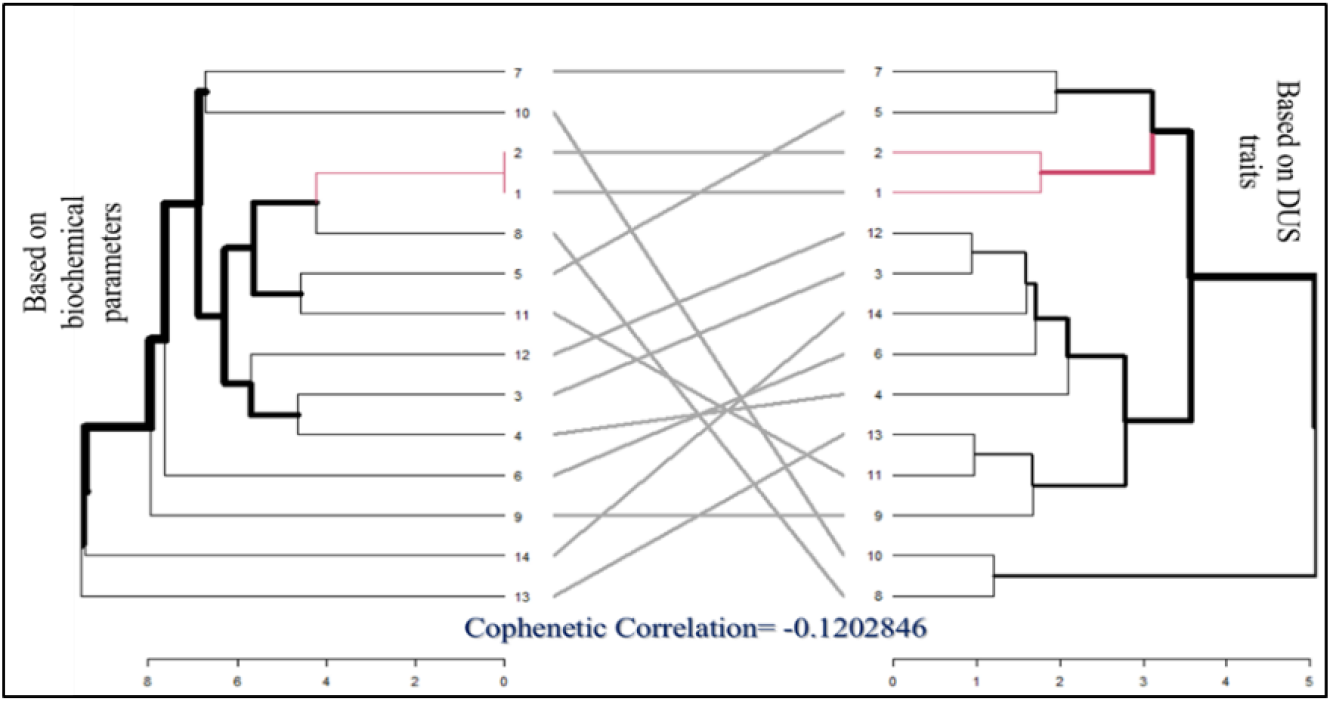
Comparative analysis between dendrogram based on biochemical and dendrogam based on qualitative traits.

The overall result suggests that there are no significant relationship among the qualitative, quantitative and biochemical characters. Plant breeders may select the genotypes for future breeding programmes with desirable characters from separate cluster of three different dendrograms.

#### Stability analysis

The AMMI model lacks a quantitative stability measure, which is necessary for quantifying and ranking genotypes in terms of yield stability (68,69). Purchase et al. (2000) (29) introduced the AMMI stability value (ASV) to measure and rank genotypes based on their yield stability. In a two-dimensional scatter gramme of IPCA1 (interaction principal component analysis axis 1) scores vs IPCA2 scores, the ASV is the distance from zero. To account for the respective contributions of IPCA1 and IPCA2 to the overall GE sum of squares, the IPCA1 score must be weighted by the proportionate difference between IPCA1 and IPCA2 scores **(Table 7)**. The Pythagorean Theorem is then used to calculate the distance from zero (Purchase et al., 2000) (29).

**Table 7.** Stability analysis of Genotypes with their IPCA1 and IPCA2 value with their corresponding ASV (Anthocyanin Content)

| Genotypes | Anthocyanin | IPCA1 | IPCA2 | ASV |
| --- | --- | --- | --- | --- |
| Cooch Behar Local 02 | 15.47643 | -1.2445 | 0.164207 | 972.6068 |
| Cooch Behar Jalpai 01 | 149.9029 | 3.579457 | 0.335963 | **7.431188** |
| **Jalpaiguri Local 02** | 16.49674 | -1.22573 | 0.144347 | 957.9342 |
| **Cooch Behar Deshi 01** | 64.21629 | 0.075914 | -0.46241 | 59.33002 |
| **Cooch Behar Local 01** | 16.1582 | -1.23202 | 0.150921 | 962.8502 |
| **K. Jyoti** | 5.551173 | -1.39323 | 0.357598 | 1088.846 |
| **Cooch Behar Badami 02** | 151.4022 | 3.672306 | 0.384449 | 2869.995 |
| **Cooch Behar Local 04** | 5.851319 | -1.38984 | 0.352047 | 1086.191 |
| **Cooch Behar Badami 01** | 94.11039 | 1.170269 | -0.45496 | 914.5935 |
| **K_Sundari** | 91.60773 | 1.072578 | -0.46712 | 838.2452 |
| **Jalpaiguri Local 03** | 29.04161 | -0.95562 | -0.08252 | 746.8401 |
| **Cooch Behar Local 03** | 17.21332 | -1.21222 | 0.130487 | 947.376 |
| **Coochbehar Deshi 02** | 62.29188 | 0.011423 | -0.45246 | 8.938532 |
| **Jalpaiguri Local 01** | 30.14652 | -0.92879 | -0.10055 | 725.8753 |

In the current experiment ASV was calculated for tuber yield and anthocyanin content in the tuber **(Table 8)**. In the ASV approach, the genotype with the lowest ASV score is the most stable; as a result, genotype 2 (Coochbehar Jalpai 01) was the most stable in terms of yield characteristics, as well as in terms of tuber anthocyanin content.

**Table 8.** Stability analysis of Genotypes with their PC1 and PC2 value with their corresponding ASV (Tuber Yield)

| **Genotypes** | **Tuber Yield** | **IPCA1** | **IPCA2** | **ASV** |
| --- | --- | --- | --- | --- |
| **Cooch Behar Local 02** | 293.6219 | -4.14345 | 1.068834 | 61.44117 |
| **Cooch Behar Jalpai 01** | 124.7864 | 0.083282 | 0.230382 | **1.256075** |
| **Jalpaiguri Local 02** | 74.91452 | 2.190084 | 0.301929 | 32.47216 |
| **Cooch Behar Deshi 01** | 93.53919 | 1.28723 | -1.0319 | 19.11269 |
| **Cooch Behar Local 01** | 73.00785 | 1.268556 | 1.065043 | 18.83808 |
| **K. Jyoti** | 105.9523 | 0.519803 | 0.140914 | 7.708023 |
| **Cooch Behar Badami 02** | 93.67484 | 0.77566 | 0.086374 | 11.50046 |
| **Cooch Behar Local 04** | 53.97773 | 1.613687 | -0.09141 | 23.92512 |
| **Cooch Behar Badami 01** | 172.2275 | 2.831107 | 1.662108 | 42.00763 |
| **K_Sundari** | 437.1598 | -8.14076 | 1.82397 | 120.7109 |
| **Jalpaiguri Local 03** | 178.1408 | 4.598436 | 2.312408 | 68.21682 |
| **Cooch Behar Local 03** | 86.45338 | 2.563262 | -5.18293 | 38.35539 |
| **Coochbehar Deshi 02** | 83.2097 | -7.02037 | -2.35997 | 104.1127 |
| **Jalpaiguri Local 01** | 61.53941 | 1.573481 | -0.02575 | 23.32885 |

## Conclusion

The results of the present investigation suggest that local potato cultivars collected from the northern part of West Bengal showed a high level of genetic variation for 30 qualitative traits with 91 phenotypic variants. In addition, association analysis among the different qualitative traits will be helpful for the plant breeder to select the preferred genotypes for further improvement. Genetic relatedness in potato cultivars revealed in this investigation can be useful for potato breeders to manage potato germplasm. Finally, the data obtained from this study will also be helpful for the DUS characterization and identification of the unique traits according to the genotypes. All the fourteen genotypes showed significant variation based on their seven quantitative characters. The differences between genotypes were highly significant at the 1% and 5% alpha level for all characters like tuber yield, length and weight of tuber, length of sprout, chlorophyll content, canopy temperature, and number of tuber per plant, according to the analysis of variance. Three quantitative traits namely tuber yield, tuber breadth, tuber length played major role in the genetic variance. Anthocyanin content had more contribution to diversify the local potato cultures according to biochemical characterization AMMI analysis suggested that Jalpai is the most stable potato cultivars in respect to anthocyanin content and tuber yield per plant.

## Acknowledgments

Authors are thankful to the Department of Science & Technology and Biotechnology (DSTBT), Government of West Bengal for funding the present Research work. We would also grateful to Director of Research, Uttar Banga Krishi Viswavidyalaya, In-Charge, Regional Research Station, Terai Zone, Uttar Banga Krishi Viswavidyalaya and In-Charge, Central Germplasm Conservation Unit, Uttar Banga Krishi Viswavidyalaya for providing the necessary plant materials and facilities for field and laboratory works during the present investigation.

## Supporting information captions

- Supplementary Table I. Estimated different allelic diversity parameters based on 29 phenotypic allele of ten DUS traits
- Supplementary Table II. ANOVA and genetic components analysis based on nine quantitative traits of potato
- Supplementary Table III. Analysis of variance among the three groups of potato revealed by DUS traits
- Supplementary Table IV. Genotypic (G) and Phenotypic (P) correlation for seven quantitative characters in Potato
- Supplementary Figure I. Dendrogram based on seven quantitative traits
- Supplementary Figure II. Contribution of seven biochemical traits towards variability using PCA

